# Combining the alpha-2 adrenergic agonist clonidine with naloxone rescues fentanyl-induced physiologic dysfunction and increases survival

**DOI:** 10.1101/2025.11.07.687263

**Authors:** Alyssa Rivera, William E Schutzer, Alex Sonneborn, Sandra L Shotwell, Aaron Janowsky, Atheir I Abbas, Randy Torralva

**Author notes:** **Address correspondence to:** Dr. Phillip Randy Torralva, TMT-RX Inc. 2603 NE 60^th^ Ave. Portland, OR 97213.

## Abstract

Fentanyl leads to tens of thousands of overdose deaths every year despite widespread availability of naloxone. Like other opioids, fentanyl causes respiratory depression. Unlike morphine, high dose fentanyl rapidly produces airway obstruction, muscle rigidity, and cardiovascular failure. Using a rat model of opioid overdose, we compared the physiological effects of fentanyl and morphine and studied the efficacy of a novel rescue strategy. In contrast to morphine, we report that fentanyl more frequently causes respiratory failure secondary to vocal cord closure and leads to more severe cardiovascular disruption, including the blockade of baroreflex-like rebound in blood pressure. We also show that administration of intramuscular naloxone immediately after intravenous infusion of fentanyl did not improve survival. However, combining intramuscular naloxone with the alpha-2 adrenergic agonist clonidine rescued vocal cord function and stabilized cardiovascular and respiratory physiology from fentanyl-induced effects. Our findings demonstrate that fentanyl is associated with a unique and more severe toxidrome compared to morphine. Also, supplementing naloxone with drugs targeting the adrenergic system improves survival primarily by reopening the upper airway, implicating airway obstruction as a significant component of fentanyl-induced respiratory depression. Therefore, reversal of vocal cord closure appears to be the necessary precursor to the restoration of not only respiration, but also vascular autoregulation, a significant determinant of survival from fentanyl overdose.

## INTRODUCTION

According to the U.S. Centers for Disease Control and Prevention (CDC), fentanyl remains the most common cause of overdose death in the U.S. and overall cause of death for U.S. adults ages 18-44^1,2^. Naloxone, the widely available mu-opioid receptor (MOR) antagonist nasal spray, has revolutionized the use of opioid overdose medications by non-medical first responders, undoubtedly saving many lives among the > 600,000 non-lethal overdoses now documented annually in the U.S^3^. However, despite recent reports of regional decreases in fentanyl-associated overdose deaths, the U.S. continues to lose > 50,000 individuals to drug overdoses per year^2^, and > 70% are due to fentanyl^1^. Ongoing research into naloxone’s efficacy against fentanyl overdose suggests that current delivery methods and formulations may not fully address the pharmacologic and physiological basis for fentanyl’s lethal toxicity^4–7^. Therefore, the present therapy and harm reduction practice of relying largely on intranasal naloxone alone or even shifting to intramuscular injection for overdose prevention, must be re-evaluated^7–9^.

Fentanyl, like all opioids, causes respiratory depression that is responsive to MOR antagonists such as naloxone or nalmefene^10,11^. However, fentanyl and fentanyl analogues (F/FAs), in contrast to morphine-based opiates, can cause atypical overdose effects that include respiratory failure from the rapid onset of muscle rigidity in the chest wall and diaphragm (wooden chest syndrome)^4^, airway obstruction from vocal cord (VC) closure (VCC)^12,13^, and severe cardiovascular and pulmonary dysfunction that can be lethal within 1-3 minutes without emergency medical support^14,15^. These atypical clinical effects are dose-dependent^16–18^, associated with limited fentanyl metabolite production (suggesting a more rapid death in < 3min) ^4,19^, and have a more limited intervention window compared to the slower onset respiratory depression seen with morphine^20,21^. Moreover, these atypical effects have been demonstrated after both clinical and illicit fentanyl administration, including in surgical settings^13^, from public health data, and at supervised injection sites^14,15^.

Experiments in both anesthetized and awake animals have similarly reported that fentanyl induces rapid full-body muscle rigidity accompanied by impairment of respiratory mechanics, apnea, hypoxemia, and death^22–25^. Interestingly, pre-treatment with alpha-1 adrenergic receptor (A1AR) antagonists^22,24^, or alpha-2 adrenergic receptor (A2AR) agonists^26,27^ is sufficient to attenuate this rigidity. This suggests that the atypical effects of fentanyl, such as VCC, may be caused by dysfunction of adrenergic signaling. We previously hypothesized that fentanyl-induced disruption of noradrenergic homeostasis is related to autonomic imbalance in brainstem nuclei controlling VC musculature, causing rapid VCC that would not be fully rescued by MOR antagonists like naloxone^5^. Importantly, to our knowledge, no studies have directly examined the effects of fentanyl and morphine on VCC, or tested the combination of naloxone with A2AR agonists as a strategy for preventing death from the atypical effects of fentanyl^6^.

Our group first demonstrated atypical upper airway effects of fentanyl in a translational animal model of overdose using fiber optic video endoscopy to monitor VC activity ^6^. We used the model to establish the potency of fentanyl and morphine to cause VCC and tested the efficacy of intravenous (IV) naloxone to reverse these effects. Surprisingly, IV naloxone failed to increase survival when administered one minute after fentanyl-induced VCC (FIVCC) at doses of 1-2mg/kg (∼25 times the maximal human equivalent dose of intranasal naloxone). Therefore, the current study compared the effects of fentanyl, morphine, and naloxone on VC function, cardiovascular, and respiratory physiology. It also evaluated the therapeutic potential of clonidine, a clinically available alpha-2 adrenergic receptor agonist, to rescue fentanyl-induced toxicity. We find that during overdose, fentanyl produces more severe cardiovascular, respiratory, and VC dysfunction compared to morphine. We also reveal that these physiological deficits, including VCC, can be rescued by supplementing naloxone with clonidine, but not by either drug alone.

## METHODS

### Drugs and Chemicals

Fentanyl and morphine were purchased through Cayman Chemicals (Ann Arbor, MI) Euthasol/pentobarbital sodium and phenytoin sodium purchased through Covetrus (Portland, ME). All other reagents were supplied by commercial sources unless noted otherwise.

### Animal Maintenance and Housing

Male rats (Charles River, Wilmington, MA) age 7-10 weeks were housed in doubles in plastic cages with free access to food in water. The housing facility was kept at a constant 21° C with lights on a 12h light:dark cycle beginning at 0600. All procedures conducted were approved by VA Portland Health Care System Institutional Animal Care and Use Committee.

### Physiological Monitoring and Surgical Techniques

On the day of experimentation, animals were weighed to allow for calculation of drug dosing, placed in individual cages, and fasted for 90 minutes prior to anesthesia. Animals were anesthetized with urethane + α-chloralose (1500 + 40 mg/kg) in 0.9% USP saline by intraperitoneal (IP) injection followed by a 90-minute induction period. Additional boluses (15% of the original dose) were administered as needed in 30-minute increments for a maximum induction time of 4 hours. Following induction, animals were immobilized atop a rodent surgical table with a homeothermic heating pad (RightTemp, Kent Scientific, Torington, CT). During the procedure, heart rate, oxygen saturation and perfusion rate were closely monitored via pulse oximetry (Physiosuite, Kent Scientific, Torington, CT).

The femoral vein was cannulated with polyethylene tubing (ID: 0.53 mm, OD: 0.965 mm, length: 43 mm, volume: 0.12 mL) for intravenous (IV) drug administration. The femoral artery was cannulated with a piezo pressure catheter (Mikro-Tip® Catheter Transducer, 2 French, Millar, Inc). Mean arterial blood pressure and heart rate were monitored via the piezo pressure catheter, recorded (PowerLab 8/35, ADInstruments – North America, CO; 1000 Hz sample rate), and analyzed (LabChart Pro. V8, ADInstruments – North America, CO). Respiratory rate was monitored via a T-type fast thermocoupled microprobe (IT-23, Physitemp Instruments, LLC, NJ) and derived from inspiration/expiration temperature differences. Thirty minutes prior to laryngeal endoscopy, glycopyrrolate (250 µg/kg, IM, hind right limb) was administered to reduce airway secretions. An oropharyngeal intubation wedge was produced from the barrel of a 1 mL plastic syringe (Air-Tite, Products Co., Inc., Virginia Beach, VA), cut to 5 cm in length and 45° angle^28^, and used to displace the tongue and epiglottis for an unobstructed view of the VCs.

To visualize the VCs, a flexible fiberoptic ureteroscope (URF-P5, Olympus, Center Valley, PA) equipped with a digital video recorder was inserted into the lumen of the wedge above the VCs for continuous observation and recording of the larynx^29^. Recordings were analyzed for time to onset and duration of VCC, which was defined as a complete closure of the glottis for a minimum of 5 seconds. If the closure persisted for 120 seconds, it was deemed sustained VCC. In addition to visualization of the VCs, a small rodent electroglottograph (EGG; Glottal Enterprises, NY) was also recorded to monitor and confirm VC functioning. To record the EGG signal, piezo-electric leads were placed on the anterior neck region on right and left sides of the thyroid cartilage/larynx. A small amount of electrical current passes through each lead, creating changes in impedance as the VCs open and close with respiration. The EGG transducer records the impedance changes, with positive deflections corresponding to open VCs and negative deflections to closed VCs, manifesting as an oscillating electrical signal on the recording. If the VCs experience sustained closure, the signal becomes flat for an extended period. The EGG was used to record data for analysis, the rate was verified and validated with the thermocouple microprobe, the VC activity was verified with laryngeal endoscopy. EGG was the preferred method for data collection as the produced wave accounts for both respiratory rate and patterns of VC activity.

### Model of Opioid Overdose

Following anesthetization and set up for physiological monitoring, rats were administered an IV bolus of multiple doses of fentanyl or morphine over 10 seconds, followed by a 0.15 mL saline (0.9%) flush. In some experiments, intramuscular rescue drug (right hind limb, naloxone and/or clonidine) was administered 15 sec after the initial fentanyl infusion. If VCC was sustained for 120 seconds, animals were euthanized. If reversal of VCC was observed, animals were monitored for 15 minutes after fentanyl administration before euthanasia.

### Statistics and Reproducibility

Data analysis in this paper was performed using a combination of GraphPad Prism (10.6.1, Boston, MA) and MATLAB (R2024a, Natick, MA). Details about the statistical and post-hoc tests used are described in the Results section and/or figure legends with the corresponding experimental findings. For slope calculation, we used the MATLAB function *polyfit* (linear) in the specified time windows. Additionally, to find the minimum value reached for each parameter throughout the paper, we used the MATLAB function *min*. For all continuous data with variability, a Bartlett’s test for homogeneity of variances was run prior to hypothesis testing to determine if parametric or non-parametric ANOVAs were appropriate. Only rats with quality recordings for all three of heart rate, mean arterial pressure, and respiration rate were used for statistics, which led to the *n* of some physiology groups being lower than their matched group for survival statistics (e.g. Figure 3C versus Figure 2A).

## RESULTS

### Monitoring the effects of fentanyl and morphine on vocal cord, cardiovascular, and respiratory physiology

Expanding on our previous clinically-relevant VCC model of fentanyl overdose^6^, the current study includes additional real-time quantification of cardiovascular and respiratory parameters. In each experiment, we recorded baseline parameters for 15 seconds followed by a ten-second infusion of IV drug or vehicle (0.9% saline). All continuous variables were transformed to represent a percentage of the mean baseline value for each animal. After drug infusion, if VCC was sustained at 120 seconds, animals were euthanized. Otherwise, physiology was recorded for a total of 900 seconds (Figure 1A).

**Figure 1.**
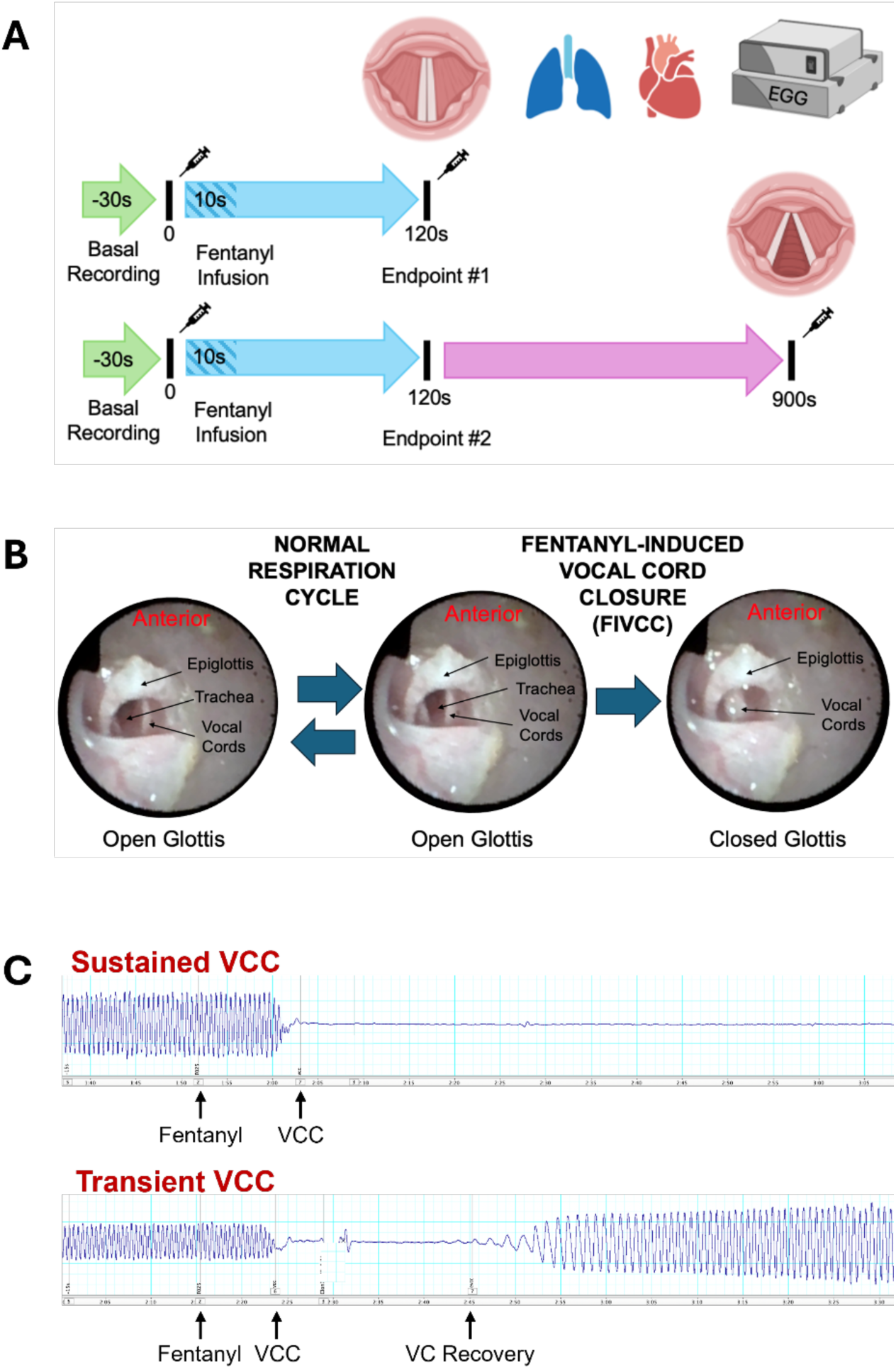
Schematic for monitoring the effects of fentanyl on cardiovascular, respiratory, and VC physiology. **A**, Animals were prepared for measurement of mean arterial pressure, heart rate, respiration rate and VC function. Baseline recordings occurred for 30 seconds (last 15 seconds used for all analyses), followed by a 10 second infusion of IV morphine, fentanyl, or saline. For animals in which drug-induced VC closure was sustained (top), the recording session ended 120 seconds after the beginning of IV infusion. For animals in which VC function was maintained or restored (bottom), the recording session continued for 900 seconds after the beginning of the infusion. **B**, Video laryngeal endoscopy demonstrates VC movement during the normal respiratory cycle and sustained fentanyl-induced VC closure (FIVCC). **C**, Electroglottogram tracings starting from −15 seconds pre-fentanyl treatment through 75 seconds post-fentanyl treatment showing sustained (top) and transient (bottom) VCC with VC recovery. Oscillations represent the opening and closing of the glottis as seen in panel **B**.

A key and novel component of the current experiments is monitoring VC activity via concurrent video laryngeal endoscopy and non-invasive EGG, during which the EGG signal is correlated with fiberoptic endoscopy as an additional measure of VC contact, movement and patency. In Figure 1B, we show example video laryngeal endoscopy photographs of typical VC functioning during normal respiration (left) and atypical VC closure after IV fentanyl infusion (right). Figure 1C depicts representative electroglottogram traces during this process showing sustained (top) versus transient (bottom) VCC after opioid administration.

### Overdose deaths caused by fentanyl and morphine occur via different mechanisms

Fentanyl and morphine were intravenously infused over ten seconds at a range of doses while monitoring VC activity. Fentanyl did not cause atypical VCC or death at the lowest dose of 0.005 mg/kg but induced sustained VCC in 100% of animals that received 0.025 mg/kg or higher (Figure 2A). Notably, almost all animals that died from fentanyl overdose experienced sustained VCC. In contrast, morphine overdose was not accompanied by sustained VCC in the majority of animals, even at the highest doses, and most of the animals died with transient VCC or no VCC (Figure 2B). For statistical analysis, we pooled deaths across all doses in each group and found that the proportion of rats that died after sustained VCC at *any* dose of fentanyl was significantly higher than the proportion of rats dying after sustained VCC at *any* morphine dose (Figure 2C, *p* = .0002, Fisher’s exact test for proportions).

**Figure 2.**
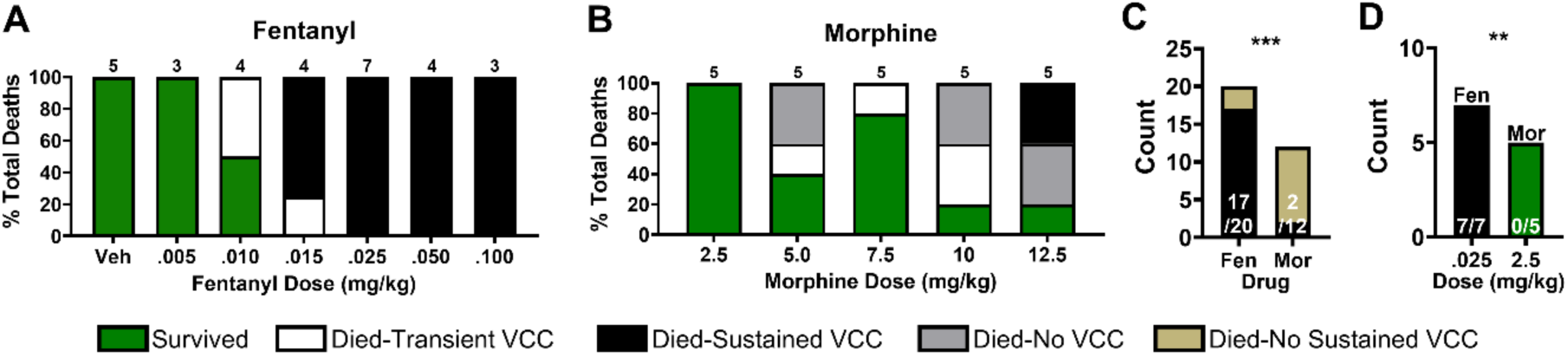
IV fentanyl induces sustained VC closure (VCC) during overdose. **A**,**B**, Percent breakdown of each cause of death in response to different doses of **A**, fentanyl or **B**, morphine. Numbers above bars represent group *n*. **C**, Pooled counts of animals that died after each drug due to sustained VCC (black) or died without sustained VCC (tan). White numbers show the proportions of sustained VCC in each group. **D**, Counts of overdose deaths at the lowest equipotent doses of fentanyl and morphine. **\*\****P* < 0.01, **\*\*\****P* < 0.001.

The MOR-dependent effects of fentanyl and morphine are considered to be equipotent at a dose ratio of 1:100 approximately^5,21^. Comparing survival after administration of our lowest equipotent doses (0.025 mg/kg fentanyl and 2.5 mg/kg morphine) revealed that fentanyl is significantly more deadly than morphine (Figure 2D, *p* = .0013, Fisher’s exact test). Since equipotent doses activate MORs to the same extent, this result suggests that fentanyl produces pharmacological activity separate from its MOR agonism. Overall, these findings indicate divergent VC physiology contributing to fentanyl versus morphine overdose, leading us to hypothesize that other critical cardiopulmonary parameters would also be mechanistically divergent between the two drugs.

### Fentanyl and morphine exert differential effects on cardiovascular and pulmonary physiology

To uncover differences in other physiological parameters during overdose, we plotted changes in heart rate (HR, Figure 3A&B), mean arterial pressure (MAP, Figure 3D&E), and respiration rate (Figure 4A&B) for each drug during the first 120 seconds after IV administration. Four fentanyl and morphine doses are shown, including three 100% lethal doses for each drug alongside a low dose with a 100% survival rate for visualization purposes. Critically, only the three 100% lethal doses (Fentanyl – 0.015, 0.025, 0.100 mg/kg; Morphine – 5.0, 10.0 12.5 mg/kg) are used for statistics as we were specifically interested in differences in overdose physiology.

**Figure 3.**
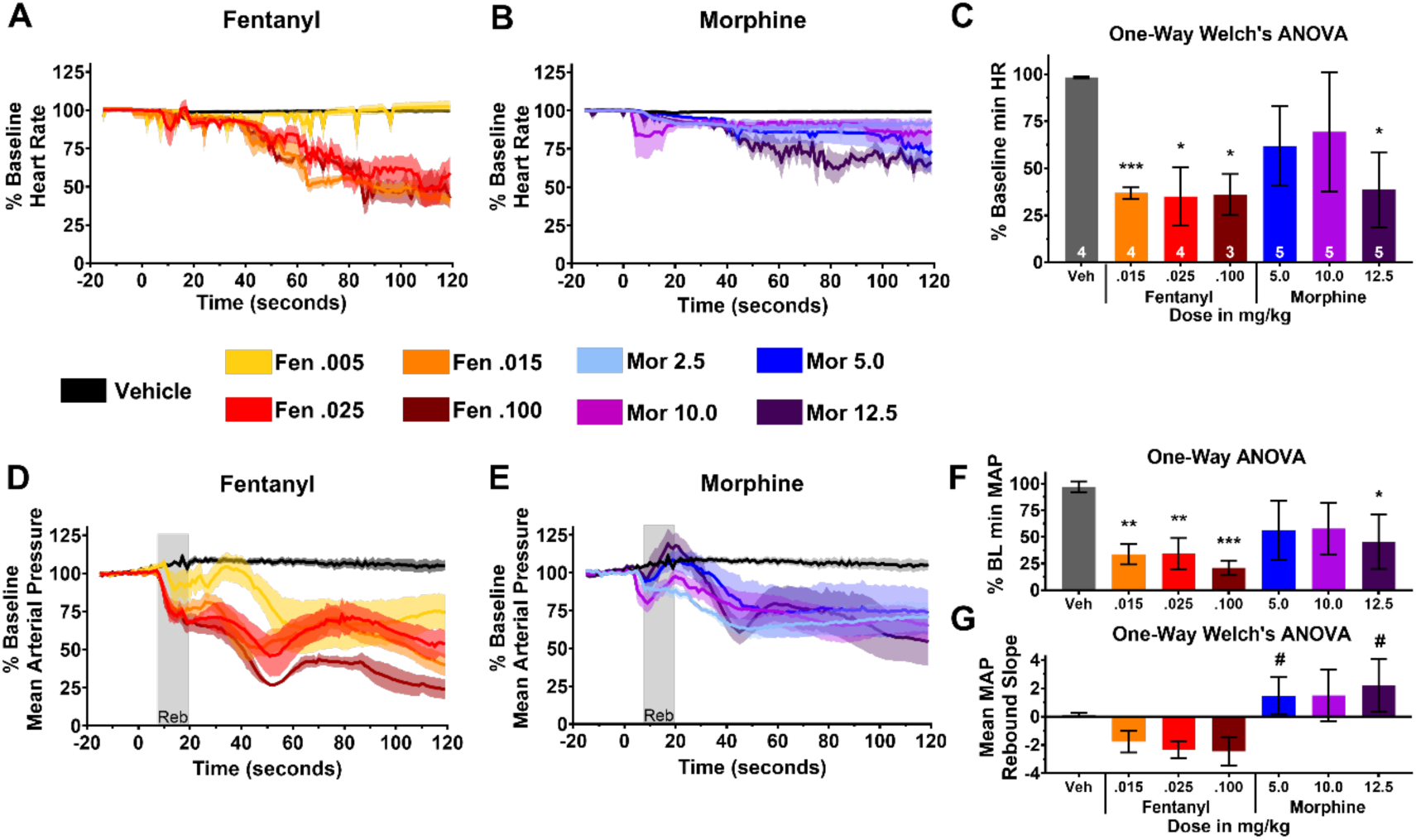
Fentanyl exerts more extreme effects than morphine on cardiovascular parameters. **A**,**B**, Time course of changes in heart rate after **A**, fentanyl or **B**, morphine administration at different doses. Means ±SEMs are shown. **C**, Bar graph of the means ±SDs of the min HR reached at each dose. Asterisks represent a significant difference compared to vehicle. White numbers inside bars indicate group *n*. **D**,**E**, & **F** Same as **A, B** & **C**, but for MAP. Note that a parametric one-way ANOVA was performed for **J** instead of a Welch’s one-way ANOVA. **G**, Bar graph of the means and SDs for slope of MAP rebound that occurs in the gray ‘Reb’ window shown in **D** or **E**. Hash symbols indicate that those morphine groups were significantly different from *all* fentanyl groups after Dunnett’s T3 post-hoc tests. Across panels, **\****P* < 0.05, **\*\****P* < 0.01, **\*\*\****P* < 0.001. Drug was applied at 0 seconds.

**Figure 4.**
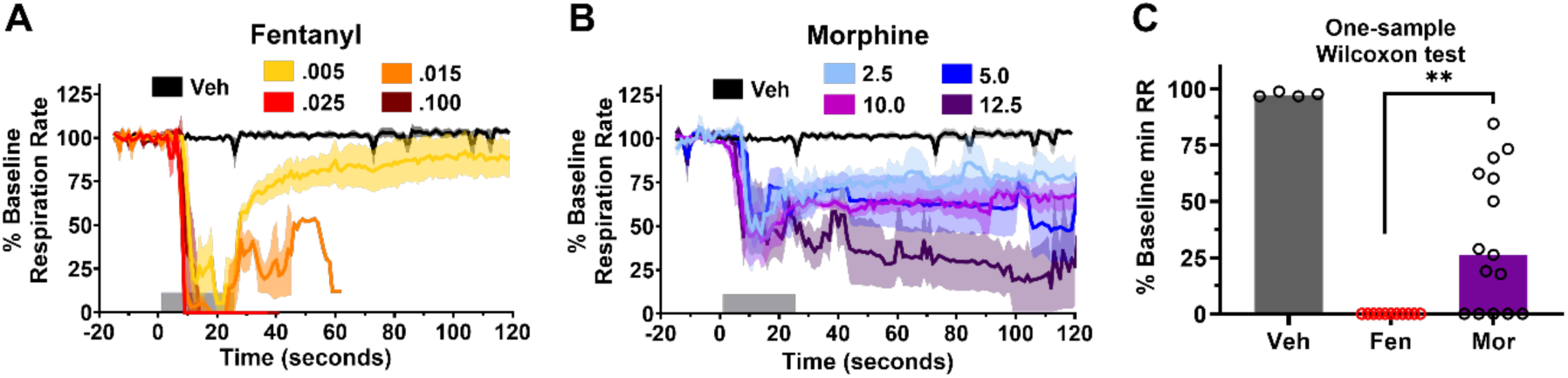
Fentanyl decreases respiration rate to a greater extent than morphine. **A**,**B**, Time course of changes in respiration rate after **A**, fentanyl or **B**, morphine administration at different doses. Means ±SEMs are shown. Gray bars at bottom of graphs indicate the time window of analysis. **C**, One-sample Wilcoxon test with Pratt correction testing if the pooled minimum RR values in the morphine group were statistically significant from zero. \*\**P* < 0.01. Drug was applied at 0 seconds.

We first asked if there were differences in the ability of fentanyl and morphine to depress cardiovascular function. To do this, we ran a one-way Welch’s ANOVA (due to unequal variances between groups, Bartlett’s test) on the mean, baseline-adjusted, *minimum* HR values reached for each dose group in the first 120 seconds after drug or vehicle administration (Figure 3C). Significant differences were found among group means (W(6, 8.408) = 204.2, *p* < 0.0001). A Dunnett’s T3 post-hoc multiple comparisons test was performed between vehicle and all drug/dose combination groups, and also between fentanyl and morphine at all doses. These post-hoc tests revealed that all fentanyl doses, and only the highest morphine dose with no equipotent fentanyl dose, significantly decreased HR (< 50%) compared to the vehicle injection (Figure 3C asterisks). Notably, the 10.0 mg/kg morphine dose, which is equipotent to the 0.100 mg/kg fentanyl dose, did not significantly reduce HR compared to vehicle (Figure 3C, magenta).

The same pattern of significance was also found with MAP (Figure 3F, F(6,23) = 5.835, *p* = 0.0008), although the statistical test used was a parametric one-way ANOVA followed by Bonferroni’s post-hoc corrections since there was no violation of equal variances. In addition, a rebound in MAP was observed shortly after morphine administration that appeared to be absent in the fentanyl group (Figure 3D&E, gray ‘Reb’ box). We quantified this rebound by fitting a slope to MAP values within the rebound time window for each animal. A Welch’s ANOVA reported a significant difference between group means (Figure 3G, W(6, 8.662) = 14.70, *p* = 0.0004). Dunnett’s T3 post-hoc tests revealed that the rebound slopes from the 5.0 and 12.5 mg/kg morphine groups were significantly higher than *all three* fentanyl groups (hash symbols, Figure 3G). Together, these data suggest that fentanyl causes more severe cardiovascular toxicity than morphine, and that the effect of fentanyl is consistent with opioid and possibly non-opioid effects that may prevent homeostatic autonomic rebound of cardiovascular function (e.g. tachycardia in response to decreased MAP/ hypotension)^30–32^.

Statistical analyses were also performed for the pulmonary parameter, respiratory rate. However, since the minimum respiratory rate reached for all overdosed fentanyl rats was zero, we could not use the same ANOVA-based analysis approach. We thus chose to test if the minimum respiratory rate reached after morphine was different than zero using a one-sample Wilcoxon test. After pooling values from all morphine rats, we found that the minimum value reached after morphine administration was indeed significantly higher than zero (Figure 4C, *p* = 0.002), leading us to reason that the effect of morphine on respiration is weaker than that of fentanyl. In fact, even at the highest doses of morphine, at no point are animals apneic in the first 120 sec, whereas animals receiving the 2 highest doses (0.025 and 0.100 mg/kg) of fentanyl had sustained apnea within 10 sec. In conjunction with the HR and MAP data, these results suggest the toxic physiological effects of fentanyl are more severe than morphine and may be due to extra-opioid pharmacological action. This led us to ask if any of these effects are due to adrenergic dysfunction (disruption of CNS noradrenergic homeostasis) that would benefit from supplementing naloxone treatment with adrenergic drugs such as clonidine.

### Combining intramuscular naloxone and clonidine improves survival by reversing vocal cord closure and stabilizing respiratory and cardiovascular function

We previously hypothesized that atypical effects of fentanyl are due to a naloxone-resistant disruption of both noradrenergic homeostasis and autonomic balance. This is consistent with our finding (Figure 3G) that fentanyl occludes/inhibits a baroreflex-like rebound in response to rapidly decreasing MAP^30^. One strategy for rescuing these effects would therefore be restoring the imbalance with adrenergic drugs. After administering 0.025 mg/kg fentanyl, the lowest dose that produced 100% death via sustained VCC (Figure 2A), we tested whether supplementing naloxone with the alpha-2 receptor agonist clonidine would improve survival. In Figure 5, a Fisher’s exact test revealed that the combination of naloxone and clonidine was indeed more effective in reducing fentanyl overdose due to sustained VCC than either naloxone or clonidine alone (*p* = .0452).

**Figure 5.**
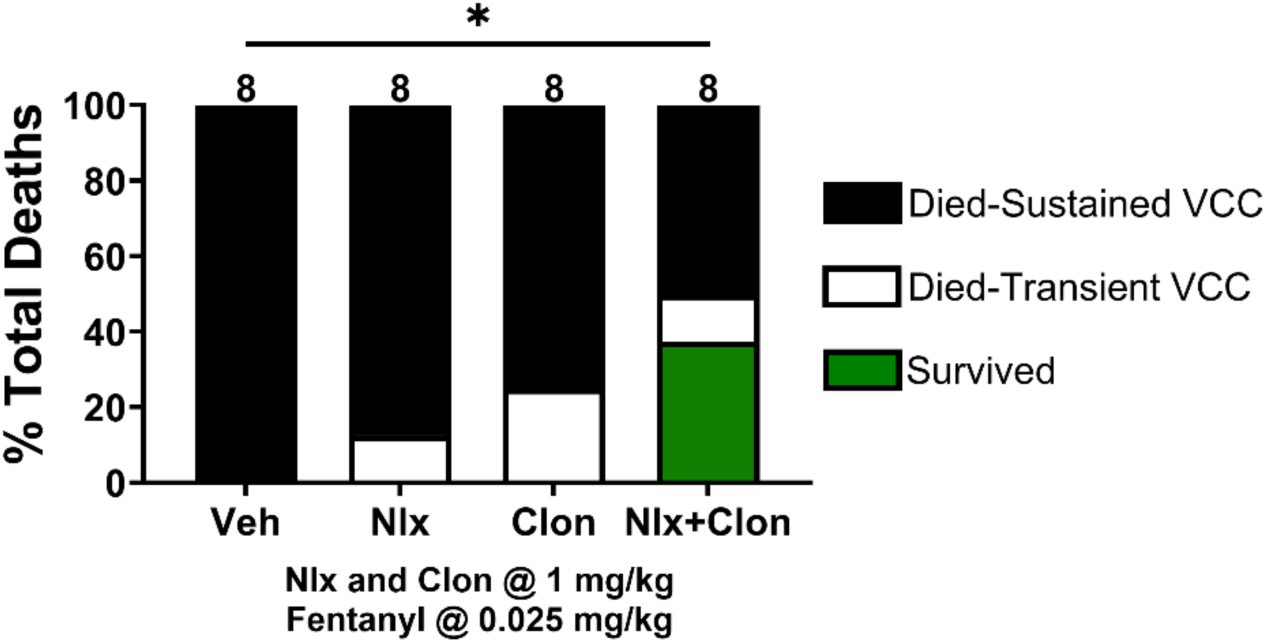
Supplementing intramuscular clonidine with naloxone increases survival after VC closure. Administration of IV fentanyl at .025 mg/kg resulted in 100% death with sustained VC closure in the vehicle group. High doses of intramuscular naloxone or clonidine alone (1 mg/kg) were not sufficient to rescue survival. Combining these doses of naloxone and clonidine rescued VC closure and caused survival in 3/8 rats. \**P* < 0.05.

We also wanted to know whether the combination of naloxone and clonidine could rescue cardiovascular parameters, such as the rebound in MAP, which would indicate a potential restoration of autonomic balance via synergy between the two drugs. We broke down the naloxone + clonidine group from Figure 5 into survivors (green, *n* = 3) and non-survivors (brown, *n* = 4, due to cardiovascular equipment failure for one rat) and plotted them alongside a control group (black, *n* = 8) where saline was injected instead of clonidine + naloxone for rescue (Figure 6A). A one-way ANOVA found significant differences between the group means of MAP rebound slope (F(2, 12) = 5.287, *P* = 0.0226), and Bonferroni’s post-hoc comparisons found that the slope of the survivor group was significantly more positive than both the control and non-survivor groups (Figure 6A, hash symbol). This result suggests that the combination of naloxone and clonidine is sufficient to restore a baroreflex-like rebound in MAP which leads to increased survival.

**Figure 6.**
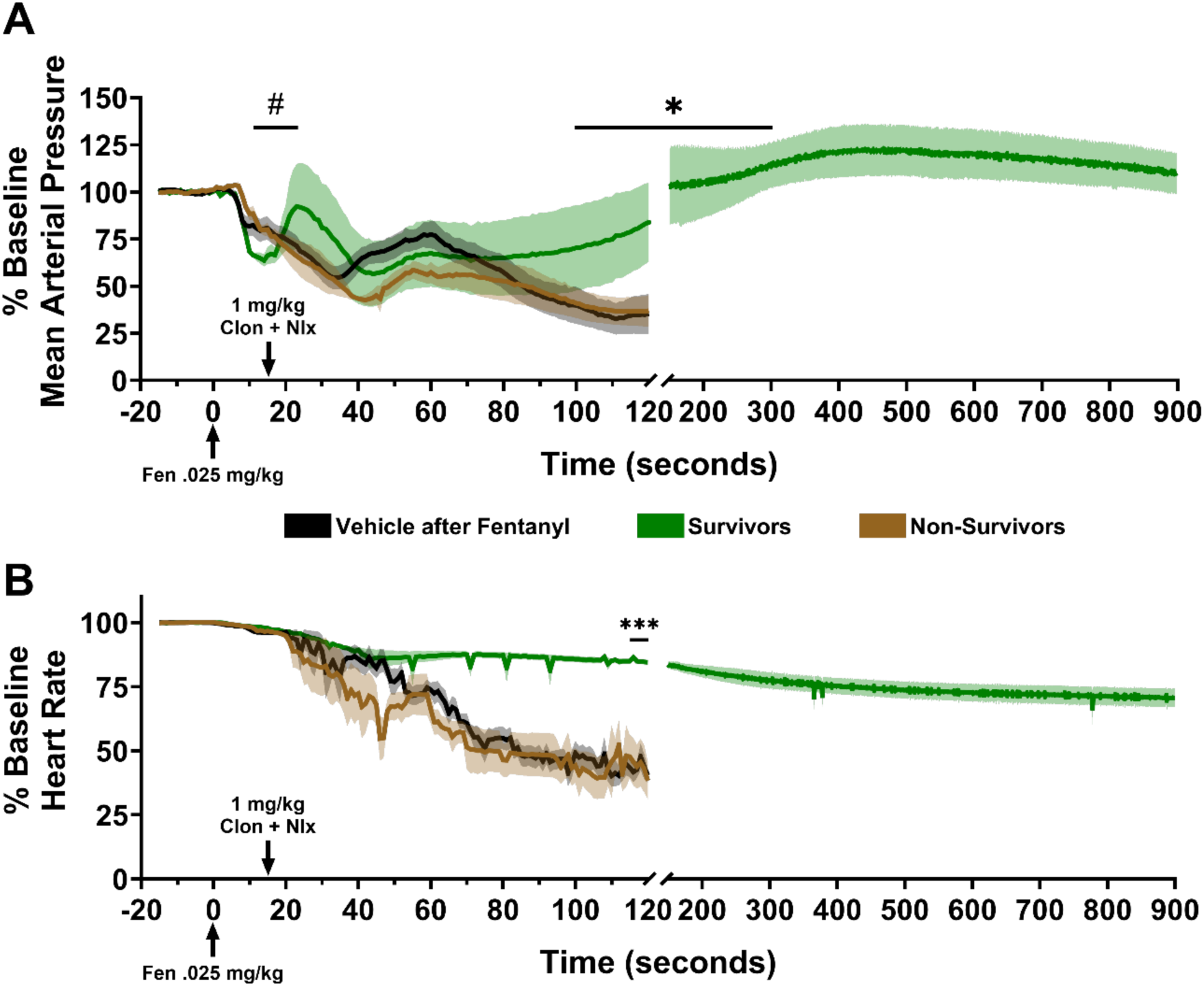
The combination of naloxone and clonidine rescues cardiovascular parameters of survivors. **A**, Baseline-corrected time courses of MAP for survivors (green) and non-survivors (brown) after Clon+Nlx rescue injection, and a control group (black) where 0.9% saline was injected instead of Clon+Nlx after IV fentanyl administration. # indicates significantly higher MAP rebound slope in survivors compared to non-survivors and vehicle groups. * represents significant MAP restoration in survivors compared to the other groups. **B**, Same as in **A**, but for heart rate. *** *P* < 0.001. Arrows indicate time of injection for the respective drugs or vehicle. Means ±SEMs are shown.

In these same animals that survived, MAP continued to increase until about 300 seconds after naloxone and clonidine administration, while continuing to decrease in the other groups until they were sacrificed at 120 seconds. It is noteworthy that MAP for all survivors remained above baseline for the duration of recording (900 sec). Using the average MAP per animal at 100-300 seconds in the survival group, and the last five seconds before death for the control and non-survival groups, a separate one-way ANOVA found significant differences between the group means of restored MAP values (F (2, 12) = 6.423, *P* = 0.0127). Bonferroni’s post-hoc tests revealed a similar pattern of significance where MAP values in the survivor group were restored to significantly higher levels than in the control or non-survivor groups (Figure 6A, asterisks). The same observation was seen for the HR parameter using a similar time window (Figure 6B, F (2, 12) = 18.39, *P* = 0.0002), with post-hoc tests finding highly significant differences between the naloxone + clonidine group and both other groups (Figure 6B, asterisks).

Finally, in Figure 7, we ran an analysis for respiration rate similar to the analysis in Figure 4. Again, since all respiration rate values in both groups of rats with no survivors (vehicle and non-survivors) went to zero, we used a one-sample Wilcoxon test to check if the median respiration rate in the survivors group returned to a value greater than zero in the naloxone and clonidine group. We found that this value was indeed significantly higher than zero over the last five seconds of recording before sacrificing animals with sustained VCC (Figure 7, *P* = 0.0068, asterisks). Taken together, these results suggest that a combination of naloxone and clonidine is significantly more effective than naloxone or clonidine alone at opening VCs and stabilizing cardiovascular and pulmonary function.

**Figure 7.**
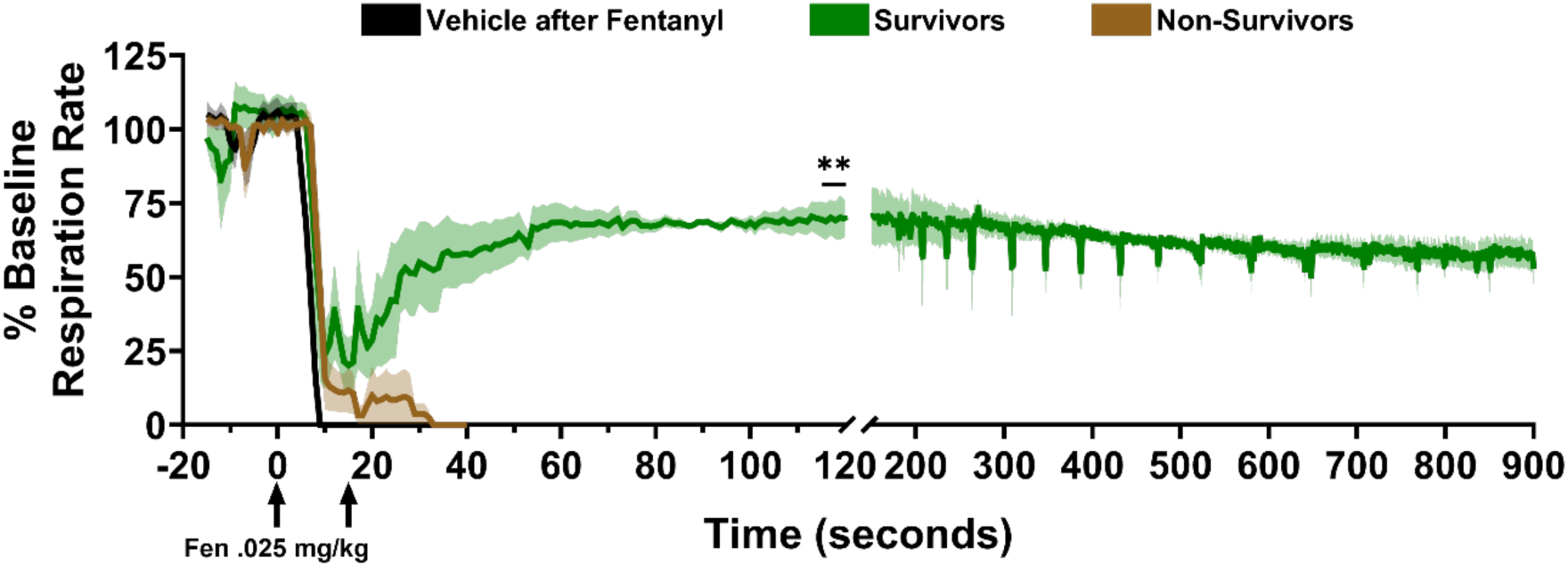
The combination of naloxone and clonidine restores respiration in survivors. Baseline-corrected time courses of respiration rate for survivors (green) and non-survivors (brown) after Clon+Nlx rescue injection, and a control group (black) where 0.9% saline was injected instead of Clon+Nlx after IV fentanyl administration. ** indicates significantly higher respiration rate than zero after one-sample Wilcoxon test. Left arrow under X-axis represents time of fentanyl injection, right arrow represents time of Clon+Nlx or vehicle injection. Means ±SEMs are shown.

## DISCUSSION

Herein, we present mechanistic evidence from an animal model^6^ of fentanyl overdose demonstrating multiple interdependent effects. These findings may explain why fentanyl deaths continue despite widespread availability of naloxone^9,14,33^. This study quantitatively demonstrates that the underlying lethal action of fentanyl is not limited to respiratory depression but involves a naloxone-resistant toxidrome that includes respiratory as well as cardiovascular effects. Our results also demonstrate a mechanistic pathway for treatment of the toxidrome: MOR antagonism combined with A2AR agonism. These findings suggest pharmacological targets for the development of enhanced opioid rescue agents and countermeasures for fentanyl weaponization^34–37^.

Our fentanyl toxidrome model (Figure 1) evaluates real-time physiological responses including HR, respiration and VC activity (Figure 2) and mirrors illicit drug use^6,38^. Bolused IV fentanyl (30mcg/kg bolused over 15 sec) induces high plasma concentrations (∼> 350ng/ml) and consistently reproduces a continuum of dose-dependent atypical clinical effects^18,39^. In this overdose model, fentanyl (25-100 μg/kg) rapidly induced lethal toxidromic effects including sustained VCC, severe bradycardia and profound hypotension in 100% of animals (Figure 2, 3). This contrasts with equipotent doses of morphine, where deaths were not attributable to VCC (Figure 2) and effects on MAP, HR and RR (Figure 3, 4) were less severe.

Importantly, these findings demonstrate fundamental differences in mechanisms of lethality between fentanyl and morphine. Fentanyl overdose more commonly causes severe hypoxemia from a mechanical failure of respiration (VCC)^12^, while morphine causes prolonged respiratory depression (Figures 2 and 4)^6,21^. It is notable that without endoscopic observation or EGG monitoring of the upper airway (e.g. studies using tracheotomized animals or plethysmography), VCC could be mistaken for apnea or respiratory depression, emphasizing the difficulty of establishing the prevalence of VCC in ongoing fentanyl deaths. Additionally, VCs and laryngeal muscles relax in animals and humans following cardiac arrest after fentanyl overdose, which precludes postmortem detection of VCC.

The predominance and consistency of FIVCC (Figure 2) is corroborated by clinical studies demonstrating its prevalence in humans^12,13^. Bennett et al., (1997), using airway endoscopy, demonstrated that > 93% of test subjects administered 20-30 µg/kg of IV fentanyl or an analogue, rapidly developed sustained VCC requiring immediate intubation to avoid hypoxemia leading to cardiac arrest^12,13^. The significance of identifying airway obstruction is non-trivial, as respiratory depression from fentanyl is responsive to IV, IM and intranasal (IN) naloxone, while VCC, as we have previously demonstrated, is not responsive to high dose IV naloxone 60 sec after VCC^6^. Thus, VCC is unlikely to be affected by slower delivery methods (e.g. IM or IN)^7,11,40^.

Physiological mechanisms for opioids involve receptor agonism in the central nervous system (CNS) that generally decreases efferent sympathetic nerve activity and increases parasympathetic (vagal) tone. However, differences between fentanyl and morphine can be explained by multiple central and peripheral opioid and non-opioid mechanisms impacting CV function. Fentanyl but not morphine^41^ antagonizes A1ARs, suggesting a mechanism for peripheral vasodilation^30,42–44^. Fentanyl also activates MORs controlling vascular tone (vasodilation) and cardiac function (decreased contractility)^45^. Additionally, it has direct relaxant effects on vascular endothelium^46^ and indirectly causes vasodilation as a response to extreme hypoxemia^47^. Fentanyl causes more extensive myocardial depressant effects than morphine, impacting contractility^16,48–50^.

Centrally, fentanyl blocks baroreceptor reflex (BRR)-mediated tachycardia that normally occurs as a response to peripheral vasodilation^30–32^. Pressure sensors in the aorta and carotids signal to cardiac vagal nuclei (CVN) in the nucleus ambiguus (NUCA) as part of the reflex arc. Fentanyl but not morphine binds A1ARs to inhibit sympathetic input to CVN. It also binds MORs that inhibit GABA inhibitory interneurons, causing bradycardia via increased parasympathetic output^31,32^. Therefore, fentanyl has a more profound effect than morphine on BRR (Figure 3), while morphine’s sympathetically-mediated MAP rebound remains intact^51^. Our CV data captures fentanyl’s ability to disrupt the primary autoregulatory BRR response to vasodilation (Figures 3D & 3E), indicating that restoration of the BRR (Figure 6A) is a critical determinant of survival, on par with restoring respiration. More specifically, the critical effect of BRR restoration may be in maintaining cardiac output (CO), optimizing coronary artery perfusion and maintaining O2 delivery to the myocardium itself. Fentanyl induced a rapid decrease in MAP and HR, suggesting a significant decline in CO with an insufficient coronary artery pressure gradient to maintain the high aerobic demands of myocardial cells^52^.

In our study, IM naloxone was ineffective at reversing VCC (Figure 5), corroborating our previous studies with IV naloxone^6^. A recent clinical study of IM versus IN naloxone evaluated treatment efficacy for fentanyl-induced respiratory depression and muscle rigidity and demonstrated IM naloxone was more efficacious and rapid in onset for reversing respiratory depression^7^. However, neither administration method consistently reversed the muscle rigidity that occurred in ∼50% of subjects and did not prevent rigidity recurrence^7^, but did prevent recurrence of respiratory depression (renarcotization) after a single dose of naloxone. The clinical continuum of dose-dependent fentanyl effects indicates muscle rigidity occurs at lower doses than VCC, suggesting that IM naloxone will not be adequate to reverse FIVCC^12,13,18,55^. Renarcotization is hypothesized to be the primary cause of ongoing fentanyl overdose deaths^53^. However, this has yet to be verified in the field or in animal studies of fentanyl doses significantly higher than those that cause rapid death from VCC^54^. The prevalence of atypical overdose presentations with fentanyl at medically supervised injection sites further supports the potential limitations of IM naloxone (see also Figure 5). Supervised injection site data show that ∼50% of illicit fentanyl overdose victims present with the rapid onset of severe muscle rigidity that is rescued by IM naloxone given immediately after overdose^14,15^. However, ∼15% of these victims present with upper airway obstruction that fails skilled airway management (positive pressure ventilation, oxygen) and is unresponsive to IM naloxone^14^. Taken together these data suggest that the majority of ongoing fentanyl deaths in the community are from a combination of 1) delayed CNS delivery of intranasal naloxone, 2) resistance of FIVCC and severe muscle rigidity to naloxone, and 3) catastrophic CV effects.

We also evaluated the efficacy of the A2AR agonist clonidine for reversing VCC (Figures 5-7). Clonidine inhibits fentanyl-induced muscle rigidity (FIMR) and therefore mitigates apnea, hypoxemia and death in anesthetized and awake animals^26,27,54,56^. These studies isolated FIMR to a disruption of noradrenergic (NA) homeostasis in the locus coeruleus (LC) and further demonstrated that pre-treatment with A1AR antagonists (prazosin), A2AR agonists and even naloxone (IV immediately after FIMR), could effectively antagonize FIMR, but studies did not examine the effects of fentanyl on VCC^24–27^. VCC likely involves a NA disruption of vagal motor neurons to laryngeal muscles that control VCs^57–60^.

Our current data using combination naloxone and clonidine (Figures 5-7) address a number of overlapping pharmacological and physiological effects that significantly impact survival in fentanyl overdose. The two most significant physiological events following combination administration were the reversal of VCC, allowing the return of respiration, and the restoration of vascular autoregulation. These events occurred in rapid sequence, with reversal of VCC preceding the CV effect (Figures 6-7). VCC reversal (Figure 5) occurs by an A2AR-mediated mechanism and is supported by the effects of other A2AR agonists we tested (e.g. dexmedetomidine-unreported observation). We hypothesize that A2ARs agonists inhibit fentanyl’s NA disruption of cholinergic innervation to vagal motor neurons, allowing for return of VC function. Conversely, A2AR antagonists increase brainstem NE release and act as respiratory accelerants at lower doses^61^, but increase rigidity induced by fentanyl and would likely worsen FIVCC^22,24,26^. Naloxone can have similar effects on VC function via cholinergic and NA mechanisms^62,63^. There is evidence that high dose naloxone may increase acetylcholine release and if given immediately after VCC, could have a similar effect as A2AR agonists^62^. However, in severe hypoxia and hypercarbic physiologic conditions, as in fentanyl overdose, naloxone increases NE release in brainstem and CNS^63^ and would be likely to worsen VCC^58,64^. In sum, naloxone + clonidine restores respiratory drive and airway patency, respectively.

Timing of the VCC reversal effect and return of vascular autoregulation suggests survival begins with reversal of VCC (restoration of gas exchange) and may be the critical first step, followed by restoration of CV autoregulation (Figure 6 & 7). Both drugs contribute to this restoration. High dose clonidine (e.g. > 0.5-1mg/kg) has two immediate independent peripheral vasoconstrictive mechanisms (e.g. A1AR agonism and A2AR-2b agonism) that directly antagonize A1AR-mediated vasodilation by fentanyl^65–67^. In turn, naloxone under hypercarbic and hypoxemic conditions (e.g., fentanyl overdose) increases adrenal epinephrine release that causes vasoconstriction, supporting vascular resistance and MAP^63^. Additionally, naloxone contributes to cardiac inotropic effects^68^. In addition to sympathetic mechanisms, clonidine restores BRR sensitivity in animals and humans by increasing parasympathetic tone in the NUCA, while naloxone has no effect^69,70^. Reversal of FIVCC and restoration of vascular autoregulation determine survival and appear to be controlled by an A2AR-mediated increase in parasympathetic tone in the NUCA. Each of these mechanisms appear to function independently in the context of BRR restoration.

## CONCLUSION

Our study suggests fentanyl interacts with non-opioid receptor targets impacting multiple physiological systems. High dose IM naloxone cannot alleviate FIVCC or the CV toxidrome on a time scale that improves survival, and higher naloxone doses may be ineffective due to CNS catecholaminergic^56,63^ and cardiopulmonary effects^71^. Restoration of vascular autoregulation is a significant event and a critical determinant of survival. Our data support the potential use of an A2AR agonist (clonidine), in combination with naloxone, for immediate reversal of the high-dose fentanyl toxidrome. Limitations of our experimental design include use of anesthetized animals and invasive airway monitoring to evaluate the entire toxidrome, complicating comparisons to awake models^21,50,72^. Nonetheless, the results are significant in that they: 1) establish the dose-dependent lethality of FIVCC, and its importance in models for therapeutics development^73,74^; 2) demonstrate FIVCC resistance to naloxone, and A2AR modulation of FIVCC and; 3) identify vascular autoregulation as critical to survival. This study suggests new pharmacological pathways for the development of reversal agents and medical countermeasures.

## ACKNOWLEDGEMENTS

National Institutes of Health, National Institute on Drug Abuse: R41DA055409 and R44DA056267 [RT]; IAA ADA12013 [AIA]. VA Senior Research Career Scientist Award RCSR-008-21S [AJ]. OHSU Physician Scientist Award Grant 60678300 [AIA]

